# A pilot assessment of avian communities and soundscapes along an Amazonian fluvial corridor

**DOI:** 10.64898/2026.05.29.728735

**Authors:** César L. Garzón-Santamaro, Mateo A. Vega-Yánez, Leonardo Vivar, Diego J. Inclán L., Pablo A. Sánchez, Mauricio G. Herrera-Madrid, Jonathan S. Valdiviezo-Rivera, Raúl Yunkar, Ángel Wasump, Mario H. Yánez-Muñoz

## Abstract

Quantifying biodiversity patterns in remote Amazonian ecosystems remains constrained by the limitations of traditional field surveys. We combined passive acoustic monitoring (PAM), machine learning, and ecoacoustic metrics to assess the taxonomic and functional structure of bird communities along a riparian gradient in the Kapawi River, Ecuadorian Amazon. A total of 2,030 recording hours were acquired using 16 Autonomous Recording Units (ARUs) deployed along a river-to-interior forest gradient (0–800 m from the riverbank). Automated detection with BirdNET yielded 92,137 records corresponding to 379 bird species. Species richness was highest at the river edge (325 species), which also harboured the greatest number of unique taxa (71 species), while interior sites showed lower but more consistent local richness. Multivariate analyses showed clear spatial segregation between riparian and interior communities. Despite this turnover, the trophic structure remained highly homogeneous (>90% similarity), dominated by insectivorous and frugivorous guilds. Generalized linear models (GLMs) indicated strong positive associations between avian species richness and key ecoacoustic metrics, with particularly pronounced effects for the Acoustic Diversity Index (ADI) and the Bioacoustic Index (BI). Spatially explicit analyses further demonstrated marked heterogeneity in acoustic structure along the fluvial gradient, reflecting fine-scale variation in soundscape composition. Together, these findings show that riparian habitats structure avian communities primarily at the taxonomic level, while functional organization remains largely conserved across the gradient. This mismatch indicates that biodiversity components respond unevenly to environmental variation, with taxonomic richness being more sensitive than functional composition. Our results underscore the potential of ecoacoustic approaches as scalable, non-invasive tools for detecting spatial patterns in biodiversity and habitat-driven community assembly in tropical systems.

## Introduction

The Amazon basin shelters the highest concentration of avian biodiversity on Earth, playing a pivotal role in global ecosystem functioning, carbon cycling, and nutrient dynamics (Terborgh et al., 1990; Remsen, 2020). Within this biome, riparian ecotones and tropical lowland forests adjacent to river systems serve as critical ecological corridors. These environments promote high beta-diversity and species turnover due to microclimatic gradients, resource pulses, and specialized habitat structures that segregate bird communities from the riverbanks to the deep interior terra firme (Naiman et al., 2005; Remsen et al., 1983). However, evaluating these intricate community assemblages in remote and logistically challenging areas like the Ecuadorian Amazon has historically been hindered by the limitations of traditional biophysical surveys (e.g., mist-netting and point counts), which suffer from observer bias, brief temporal windows, and low detection rates for rare or cryptic species (Robinson et al., 2018). Likewise, traditional observations often overlook important components of biodiversity, such as population dynamics and fine-scale temporal changes, which are essential to measure given the rapid transformation of Amazonian ecosystems. In response to this need, tools capable of collecting biodiversity data in a rapid, continuous, standardized, and large-scale manner have emerged, with passive acoustic monitoring (PAM) standing out as a powerful alternative (Gibb et al., 2019; Alcocer et al., 2022). Over the last decade, passive acoustic monitoring (PAM) has emerged as a transformative tool in conservation biology and landscape ecology, offering continuous, non-invasive, and standardized temporal sampling across extensive spatial scales (Sugai et al., 2019; Deichmann et al., 2017). When combined with modern Artificial Intelligence (AI) algorithms, such as convolutional neural network frameworks (e.g., BirdNet architectures), PAM enables the processing of massive audio datasets, bypassing the traditional bottleneck of manual sound spectrogram analysis and providing highly reliable species-level identification (Priyadarshani et al., 2018; Wood et al., 2021). Furthermore, the calculation of multi-metric acoustic indices such as the Acoustic Complexity Index (ACI) and the Normalized Difference Soundscape Index (NDSI) allows ecologists to capture structural properties of the soundscape, partitioning biological choruses (biophony) from anthropogenic stressors (anthropophony), such as river vessel traffic (Depraetere et al., 2012). Despite these technological advancements, a critical gap persists in ecoacoustic research: most studies utilize acoustic data merely to compile taxonomic checklists or estimate raw species richness (alpha diversity), leaving the functional dimensions of the soundscape largely unexplored (Toledo et al., 2023). Ecosystem stability and resilience do not depend solely on taxonomic counts, but on functional structure and dietary guild redundancy, which dictate how communities exploit niche space and maintain ecosystem services like seed dispersal, pollination, and trophic pest control (Sekercioğlu et al., 2004; Bregman et al., 2016). In tropical riparian systems, analyzing whether high taxonomic turnover across environmental gradients translates into functional divergence or if, conversely, communities exhibit high functional redundancy despite species replacement remains an open question essential for understanding ecosystem assembly rules. In this study, we provide a comprehensive, multi-dimensional assessment of avian community composition and soundscape structure along the Kapawi River (or Capahuari River) in the Wayusentza community, Pastaza Province, in the Ecuadorian Amazon. Using an intensive passive acoustic monitoring (PAM) design, we collected 2,030 hours of continuous recordings across terrestrial gradients, standardized through rigorous machine-learning validation. We tested how avian taxonomic richness and functional dietary guilds vary with distance from the river and across spatial replicates. We hypothesized that the riparian edge (0 m) functions as a high-productivity ecotone supporting elevated species richness and specialized taxa, whereas interior forests maintain functional integrity through high trophic guild redundancy. By integrating AI-driven bioacoustics with functional trait ecology, our approach establishes a scalable framework for monitoring Indigenous-managed Amazonian landscapes and for quantifying the ecological footprint of emerging anthropogenic soundscapes in otherwise intact Neotropical systems.

## Materials and Methods

### Area of study

The Achuar community of Wayusentza is located in the rural parish of Montalvo, Arajuno Canton, in the south-central Ecuadorian Amazon, within the ancestral territory of the Achuar Nation of Ecuador. It is situated in the middle Pastaza River basin, at approximately 2°26′ S and 76°55′ W, at an elevation of 230–260 m above sea level (Fig. 1). The landscape consists of an extensive Amazonian alluvial plain with predominantly flat to gently undulating topography and slopes of less than 5%, interrupted by fluvial terraces, paleochannels, and small seasonally flooded depressions. The hydrographic network is dominated by the Pastaza River and numerous tributaries of both whitewater systems, including streams and seasonal oxbow lakes that regulate the hydrological dynamics of the landscape. The region has a warm humid climate, with annual precipitation exceeding 3,000 mm and mean temperatures ranging from 24 to 26 °C. Vegetation corresponds to the lowland evergreen rainforest of the Amazon (MAE, 2013), characterized by primary forests with a continuous canopy reaching 30–40 m in height, high floristic diversity, abundant palms, and traditional agroforestry plots managed by the community. Wayusentza is home to approximately 200 inhabitants and represents a biocultural landscape of high importance for the conservation of biodiversity and the preservation of the ancestral ecological knowledge of the Achuar people.

**Figure 1.**
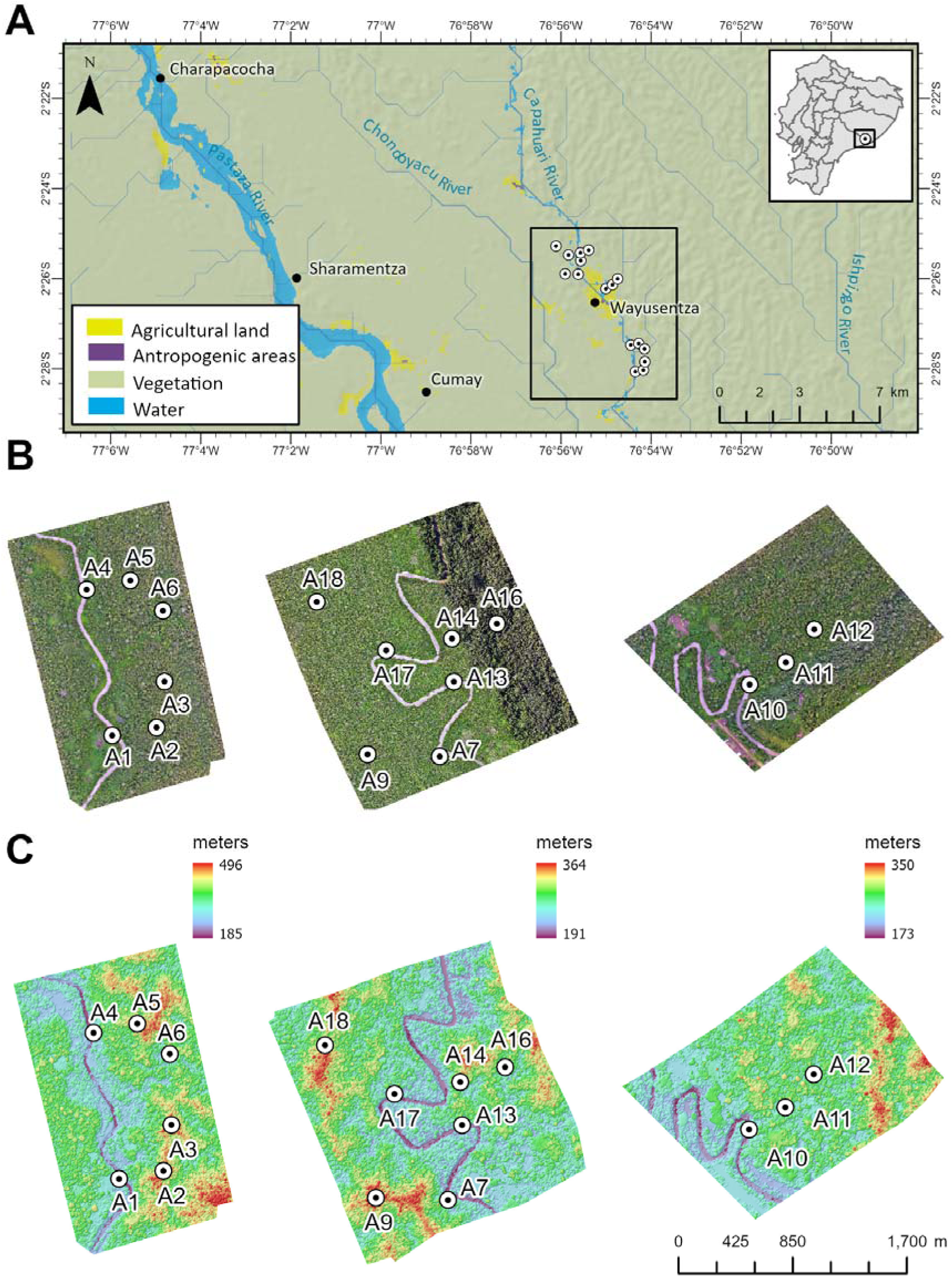
**A**. Location of the Wayusentza community in the Ecuadorian Amazon. **B.** Drone images of the study area. **C.** Digital Surface Model (DSM) of the study area. Sampling units (SU) were defined as spatial clusters of autonomous acoustic recorders (AudioMoths), distributed as follows: SU1 (A1, A2, A3), SU2 (A4, A5, A6), SU3 (A7, A9), SU4 (A10, A11, A12), SU5 (A13, A14, A16), and SU6 (A17, A18).

### Experimental design

Data for this study were collected using 16 bioacoustic recorders, stratified across six sampling units along the Kapawi River. Each sampling unit included three recording points to assess sound attenuation as a function of distance from the river: 20–25 m from the riverbank, 400 m inland, and 800 m inland. At each sampling point, an Audiomoth (Autonomous Recording Units) was installed at a height not exceeding 2 m. This device is an autonomous acoustic recorder equipped with a wide-frequency, omnidirectional MEMS microphone (Hill et al., 2018). For this study, the units were configured to record WAV files with a bandwidth of up to 48 kHz, using a duty cycle of 1 minute of recording followed by 10 minutes of silence, continuously over a 24-hour period. This schedule was selected to effectively capture daily bird activity, including crepuscular, diurnal, and nocturnal species. Recordings were conducted from March 28 to April 8, 2026.

### Bioacoustic analysis

Recordings across sampling units generated a substantial volume of data, ensuring strong spatial and temporal representativeness. In total, 132,845 WAV audio files (723.27 GB) were collected, corresponding to 2,030 hours of continuous recordings (approximately 84.6 days). The dataset was structured according to the original sampling frequencies (4, 8, 16, and 24 kHz) to enable targeted analyses of biophonic and anthropophonic components within the audible spectrum. To promote transparency and reproducibility, all derived acoustic indices, raw WAV recordings, and segmented bird vocalizations have been made publicly available in open-access repositories on Figshare, including the acoustic indices dataset (DOI: 10.6084/m9.figshare.32493456) and the WAV files and bird song segments dataset (DOI: 10.6084/m9.figshare.32494023). All files were standardized using a consistent naming convention (AM#KAPW2_AAAAMMDD_HHMMSS.wav), incorporating recorder identity, date, and time of acquisition. Pre-processing included amplitude normalization and, where required, signal resampling. Bird vocalization detection was conducted using a Python-based pipeline built on BirdNET 2.4 (Kahl et al., 2021), alongside the libraries pandas, datetime, soundfile, librosa, numpy, and scipy. No additional transformations were applied, as signals exhibited no substantial variation in recording quality or parameters. The model was run using site-specific geographic coordinates, the corresponding week of the year for each recording, and a minimum confidence threshold of 0.25.

Analyses were performed without segment overlap, using a sensitivity of 0.5, a segment duration of 3 seconds, and a frequency range of 0–15 kHz. Audio speed was fixed at 1, and FFT parameters were set to 5.33 ms. Subsequently, audio segments were extracted based on the 0.25 confidence threshold, generating up to 100 segments per file, using parallel processing (14 threads) and batches of 300 files.

Outputs were exported in Comma-Separated Values format, including the original file name, start and end times of each detection, taxonomic identity (scientific and common names), and associated confidence levels. Anthropophony detection was conducted using a classification model based on Perch v2 (van Merriënboer et al., 2025), implemented in Python with pandas, soundfile, librosa, numpy, and scipy. The analysis focused on frequencies between 1 and 10 kHz, with particular emphasis on the 1–2 kHz range associated with mechanical noise sources. FFT window sizes of 1024 and 2048 points were used. Classification into geophony, biophony, and anthropophony relied on pre-trained models; therefore, outputs were validated by an ornithologist to account for potential label uncertainty. The soundscape was characterized using the Normalized Difference Soundscape Index (NDSI; Kasten et al., 2012), defined as (β − α)/(β + α), where β represents acoustic energy in the biological band (2–8 kHz) and α corresponds to energy in the anthropogenic band (1–2 kHz). The index was computed using a 1024-point FFT and applied exclusively to time intervals with relevant detections. Additionally, sound pressure level (SPL) was estimated using Python scripts with scikit-maad, pandas, tqdm, joblib, and soundfile. Analyses were conducted across the full spectrum associated with motor detections, with emphasis on octave and third-octave bands within the 1–2 kHz range, following standard environmental noise assessment protocols. Finally, multiple acoustic indices were calculated, including the Acoustic Complexity Index (ACI; Pieretti, Farina & Morri, 2011), Acoustic Diversity Index (ADI), Acoustic Evenness Index (AEI; Pijanowski et al., 2011), and Bioacoustic Index (BI; Boelman et al., 2007), using numpy, soundfile, librosa, scipy, multiprocessing, and scikit-maad. ACI was computed using a 512-point FFT across the full spectrum up to the Nyquist frequency. ADI and AEI were estimated within the 0–10 kHz range using 1 kHz bins and a −50 dB threshold, while BI was calculated over the 2–8 kHz range with a 512-point FFT. All analyses were implemented in Python through automated scripts, ensuring workflow reproducibility and consistency in acoustic data processing.

### Statistical analysis

All analyses were conducted in R (R Core Team, 2025). An initial exploratory analysis of the previously collected data was performed, including species richness across recording stations and the variation of acoustic indices among stations. This step aimed to develop a more comprehensive understanding of the soundscape structure and its relationship with the bird community in the study area, facilitating the identification of relevant spatial and ecological patterns prior to formal modelling. Acoustic indices (ACI, ADI, AEI, BI, NDSI, and SH) were standardized using z-score transformation to place predictors on a common scale and allow comparison of effect sizes across variables. This standardization ensured that regression coefficients could be interpreted as comparable measures of relative influence on species richness. Species richness was treated as a count response variable. To evaluate the relationship between acoustic indices and avian species richness in the study area, we fitted a generalized linear model with a negative binomial error distribution using the MASS package. This approach was used to explore the ecological and bioacoustic relationships between soundscape structure and bird diversity (Sueur et al., 2008; Ross et al., 2021). The negative binomial framework was selected over a Poisson model due to clear evidence of overdispersion in the response variable, where the observed variance substantially exceeded the mean, violating a core assumption of Poisson regression (Hilbe, 2011). Overdispersion was evaluated through inspection of model residual deviance and dispersion ratios (Ismail & Jemain, 2007). In contrast to the Poisson model, the negative binomial formulation incorporates an additional dispersion parameter, allowing variance to increase independently of the mean and thereby providing more robust standard error estimation and more reliable inference under heterogeneous ecological conditions. This modelling framework allowed the estimation of standardized effect sizes for each acoustic index, enabling direct comparison of their relative contributions to patterns of species richness and providing ecological insight into how soundscape properties reflect avian |biodiversity in the study system (Ross et al., 2021).

### Drone image processing

During the field campaigns, geospatial information of the study area was collected using a DJI Mavic 3M drone. The images acquired were subsequently processed using Agisoft Metashape Professional v.1.5.2, with the project organized into three separate chunks. A total of 2,509 aerial photographs were processed, achieving an approximate alignment efficiency of 99.8%. During the alignment stage, an average of 974,799 corresponding points per chunk were generated, which allowed for establishing the spatial correspondence between overlapping images and reconstructing the initial geometry of the photogrammetric model. Subsequently, medium-quality depth maps with light filtering were generated, from which dense point clouds were constructed with an average of 138,929,355 points per chunk. Based on these products, the digital elevation models and final orthomosaics were generated. The Digital Surface Models (DSMs) achieved an average spatial resolution of 23.8 cm/pixel, while the orthomosaics had an average spatial resolution of 5.95 cm/pixel. This resolution enabled the production of high-spatial-resolution cartographic products suitable for visual interpretation, spatial analysis, and detailed characterization of the study area.

### Spatial distribution of acoustic indices

The spatial analysis was performed in R, using the sf package for vector data management and the gstat package for spatial interpolation using the Inverse Distance Weighting (IDW) method (Wong, 2016). Based on the sampling points, a regular grid with a spatial resolution of 100 meters was generated, used as a basis for the continuous estimation of acoustic indices. IDW interpolation was applied independently for each acoustic index (SH, NDSI, ACI, ADI, AEI, and BI), all derived from the processing of acoustic records. This method assigns higher weight to the observations closest in space, with a decreasing contribution as a function of distance, under the assumption of spatial autocorrelation (Isaaks & Srivastava, 1989). The procedure allowed for the generation of continuous surfaces for each index, facilitating the characterization and comparison of spatial patterns of acoustic activity within the study area.

### Spatial distribution of acoustic indices

To assess the structure and spatiotemporal variation of the bird community along the Kapawi River, acoustic data were analyzed using a taxonomic and functional (trophic groups) approach, employing parametric and multivariate statistical methods in R and PAST (Hammer et al. 2001). Initially, alpha diversity was quantified using total cumulative species richness and the count of species exclusive to each station (Sampling Units 1 through 6) and distance bands perpendicular to the river channel (0 m, 400 m, and 800 m). To determine whether there were statistically significant variations in average local species richness along the distance gradient, a one-way, one-tailed analysis of variance (ANOVA) was applied, supplemented with Welch’s test to correct for potential heteroscedasticity, having previously validated the assumptions of homogeneity of variances and normality using Levene’s and Shapiro-Wilk tests, respectively. The functional structure was determined by taxonomically classifying birds into nine distinct foraging groups based on their foraging ecology. Biocenotic affinities and community turnover were assessed using multivariate analyses based on distance matrices and similarity indices. Specifically, hierarchical clustering dendrograms were constructed using the unweighted pair group method with arithmetic mean (UPGMA) to identify basal ecological discontinuities. Finally, spatial arrangements were performed using two-dimensional non-metric multidimensional scaling (MDS); in the functional models, the coordinate vectors corresponding to each trophic trait were projected onto the bioacoustic biplot to quantitatively identify which feeding niches exerted the greatest ecological weight or control on the trophic segregation of avifauna along the monitored river system. All hypothesis tests used a significance level of α = 0.05.

## Results

### Distribution of absolute wealth across the sampling units

Through passive acoustic monitoring, a total of 92,137 detections with a confidence level of ≥45% were recorded, corresponding to 379 bird species detected by the 16 Audiomoth devices across the six sampling units (SU) assessed along the Kapawi River (Appendix 1). The absolute species richness per unit ranged from 203 to 279 (x = 236.2 species/SU). The highest species richnesses were concentrated south of the Wayusentza community settlement, in SU 2 (*n* = 279 spp.) and SU 1 (*n* = 276 species) (Fig. 2A), followed by SU4 (*n* = 235 spp.) and SU5 (*n*= 215 spp.), to the north of the village. The lowest values of absolute species richness were recorded in SU3 and SU6, with 209 and 203 spp., respectively (Fig. 2A). The number of unique species (exclusive to a single unit) showed a heterogeneous pattern. SU2 exhibited, by a significant margin, the highest number of exclusive species (*n*= 24 spp.), followed by SU1 (*n*=15 spp.) located south of the town center (Fig. 2A). In contrast, the remaining units had a reduced number of unique species, ranging from a minimum of four species in SU5 to a maximum of eight species in SU3 and SU4, all located north of the town center.

**Figure 2.**
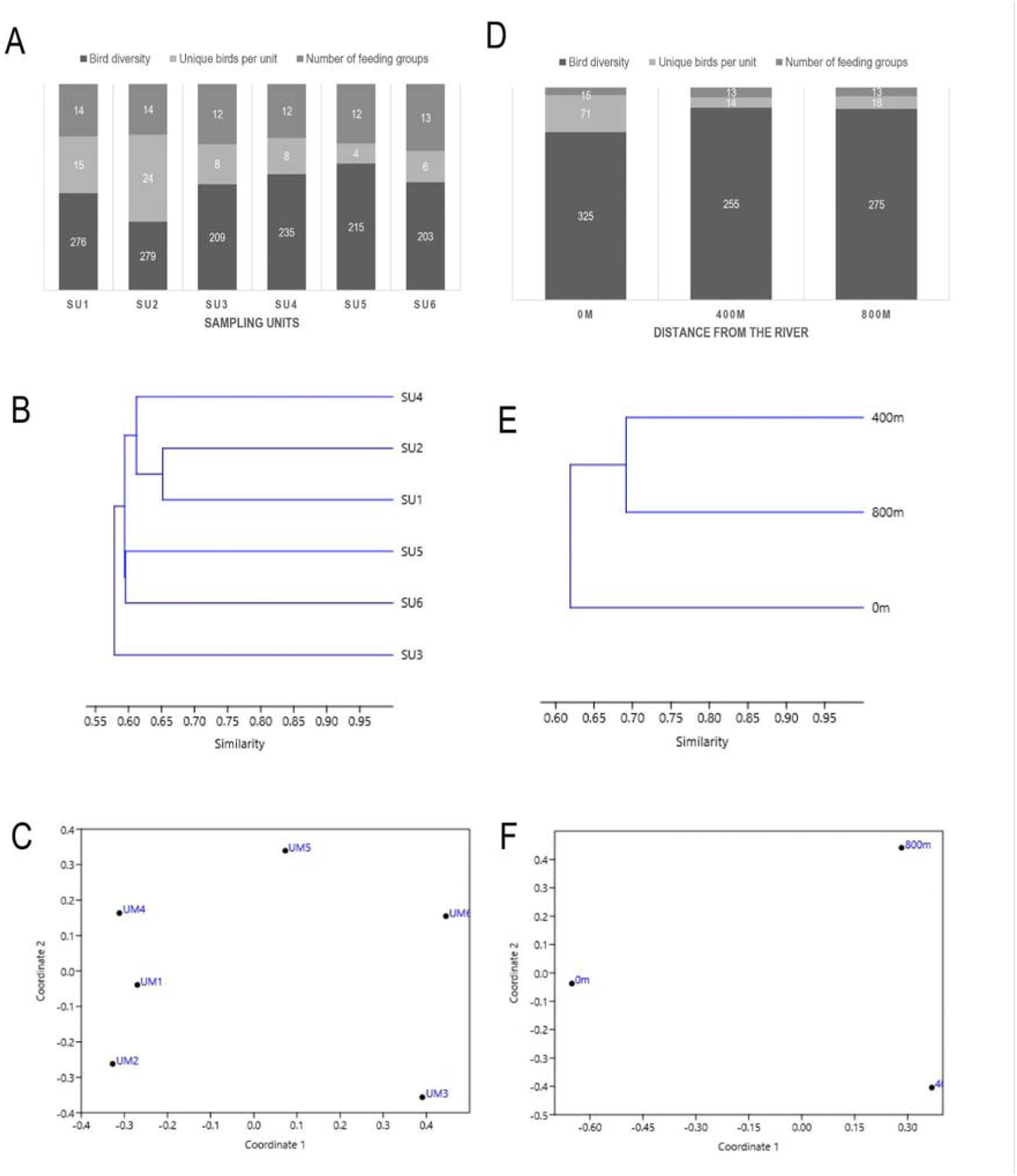
Distribution of bird species richness in the Kapawi River, Wayusentza community. (A) Distribution of species richness, number of unique species, and number of trophic groups per sampling unit; (B) Cluster analysis based on the Jaccard algorithm for six sampling units; (C) MDS analysis of presence-absence data for six sampling units; (D) Distribution of species richness, number of unique species, and number of trophic groups by distance gradient; (E) Cluster analysis based on the Jaccard algorithm for three distance gradients; (F) MDS analysis of bird presence-absence data across three distance gradients.

### Similarity and Spatial Ordering (MDS) of Absolute Wealth: Sampling Units

Cluster analysis based on the Jaccard similarity index showed that the bird communities of the six units share a basal similarity of over 55% (Fig. 2B). A clearly defined main group was identified, consisting of units SU1 and SU2, which shared the highest percentage of biological similarity in the study (65%). UM 4 was externally associated with this pair of units, with an approximate similarity of 61%. On the other hand, units SU5 and SU6 formed a second group with a similarity close to 60%. Finally, SU3 was the most dissimilar and segregated unit in the set, linking to the rest of the system below the 58% similarity threshold.

Non-metric multidimensional scaling (MDS), used to assess the spatial distribution patterns of bird species among the monitored stations, showed a clear segregation and clustering of the sampling units along the river system (Fig. 2C). The two-dimensional configuration divides the study area into three well-defined blocks or gradients based on the structure and richness of their communities:

#### High-Richness Cluster

A compact cluster was identified in the left quadrant of the biplot (Fig. 2), consisting of SU1, SU2, and SU4. These three stations exhibit a strong biological affinity with one another, which directly corresponds to the highest peaks of total taxonomic richness and trophic group richness recorded in the study.

#### Transition Gradient

SU5 was positioned in isolation in the upper-central region of the ordination (Fig. 2), acting as a turning point or intermediate ecotone between the stations with maximum biodiversity and the river sections with lower species representation.

#### The Taxonomic Contraction Block

SU6 and SU3 are located at the far right of the ordination. Although they are at opposite ends of Axis 2, indicating a divergence in the composition of their specific species, both are clearly separated from the main cluster due to their lower overall richness. This spatial pattern confirms that the bird community of the Kapawi River is not distributed uniformly, but rather exhibits marked spatial variations in which geographical extremes or local disturbances may influence the observed taxonomic richness.

### Distribution of absolute species richness along the distance gradient from the river

Bird species diversity as a function of distance from the Kapawi River bank (0 m, 400 m, and 800 m) showed marked variations in taxonomic richness and species exclusivity (Fig. 2D), with an average absolute richness of x = 285 spp./m and x = 33 unique spp./m.

The greatest species richness was concentrated directly on the riverbank (0 m), with a total of 325 species recorded. As one moved inland into the terrestrial ecosystem, species richness decreased (*n* = 255 spp.) in the unit located at 400 m, and then increased slightly to *n* = 275 spp. at the most distant point evaluated (800 m). The number of unique species (exclusive to a single distance) showed a markedly asymmetric distribution pattern. The riparian zone (0 m) exhibited the greatest biological uniqueness with *n* = 71 exclusive species (Fig. 2D). In contrast, the inland units showed considerably lower values, reaching *n* = 14 unique species in the unit at 400 m and *n* = 18 exclusive species in the unit at 800 m.

Cluster analysis based on the Jaccard algorithm showed that the three distances share a basal biological similarity of over 60% (Fig. 2E), with a clear affinity between the two forest interior sites (400 m and 800 m), which share 69% of their bird composition. In contrast, the riparian vegetation (0 m) was segregated from this main group, clustering externally with a lower similarity (62%) (Fig. 2E).

Despite these notable differences in total biodiversity aggregates, analysis of variance (ANOVA) showed that the average species richness per sampling point does not vary in a statistically significant way among the three distances (*F_5,12_*=1.012, *p* = 0.4523, Test de Welch: *p* = 0.7236). The internal biological variation within each distance was considerable: The shorelines (0 m) averaged x =158.17 ± 47.9 spp., showing high variability due to local asymmetries (maximums of *n* = 209 spp. in UM2 versus minimums of *n* =100 spp. in UM4). The intermediate zones (400 m) recorded the lowest average (x =127.67 ± 62.9 spp.), heavily penalized by the complete absence of detections (“0”) in UNIT 3.

The forest interior (800 m) exhibited the greatest consistency and the highest average local species richness (x =151.0 ± 41.5 spp.), reaching its peak in UNIT 6 with *n* = 233 species. The assumptions of normality for these models were validated using the Shapiro-Wilk test, confirming normal distributions in the stable units (*p* > 0.05), with the sole exception of UNIT 6 (*p*= 0.021) due to its strong skew toward high values in the interior.

This radical separation was clearly confirmed by non-metric multidimensional scaling (MDS). Axis 1 of the MDS (Coordinate 1), which explains the largest proportion of biological variance, segregated the riparian zone from the mature forest in a polarized manner (Fig. 2F). The results confirm the existence of a strong hydrodynamic effect in the Kapawi River, as distance from the channel modulates the composition and turnover of avifauna in the landscape, isolating the riparian zone from the dynamics of the interior understory.

### Functional structure and composition of trophic groups in the avian community

The functional structure, as represented by the number of trophic groups, remained highly homogeneous across the sampling gradient. SU1 and SU2 harbored the greatest trophic diversity (*n =* 14 trophic levels), closely followed by SU6 (*n =* 13 trophic levels). The remaining units (SU3, SU4, and SU5) recorded a constant composition of *n* = 12 trophic levels. Analysis of dietary functional traits showed a highly specialized taxonomic structure dominated overall by insectivorous species (Ins), followed in order of abundance by frugivorous (Fru) and carnivorous (Car) species (Fig. 3A).

**Figure 3.**
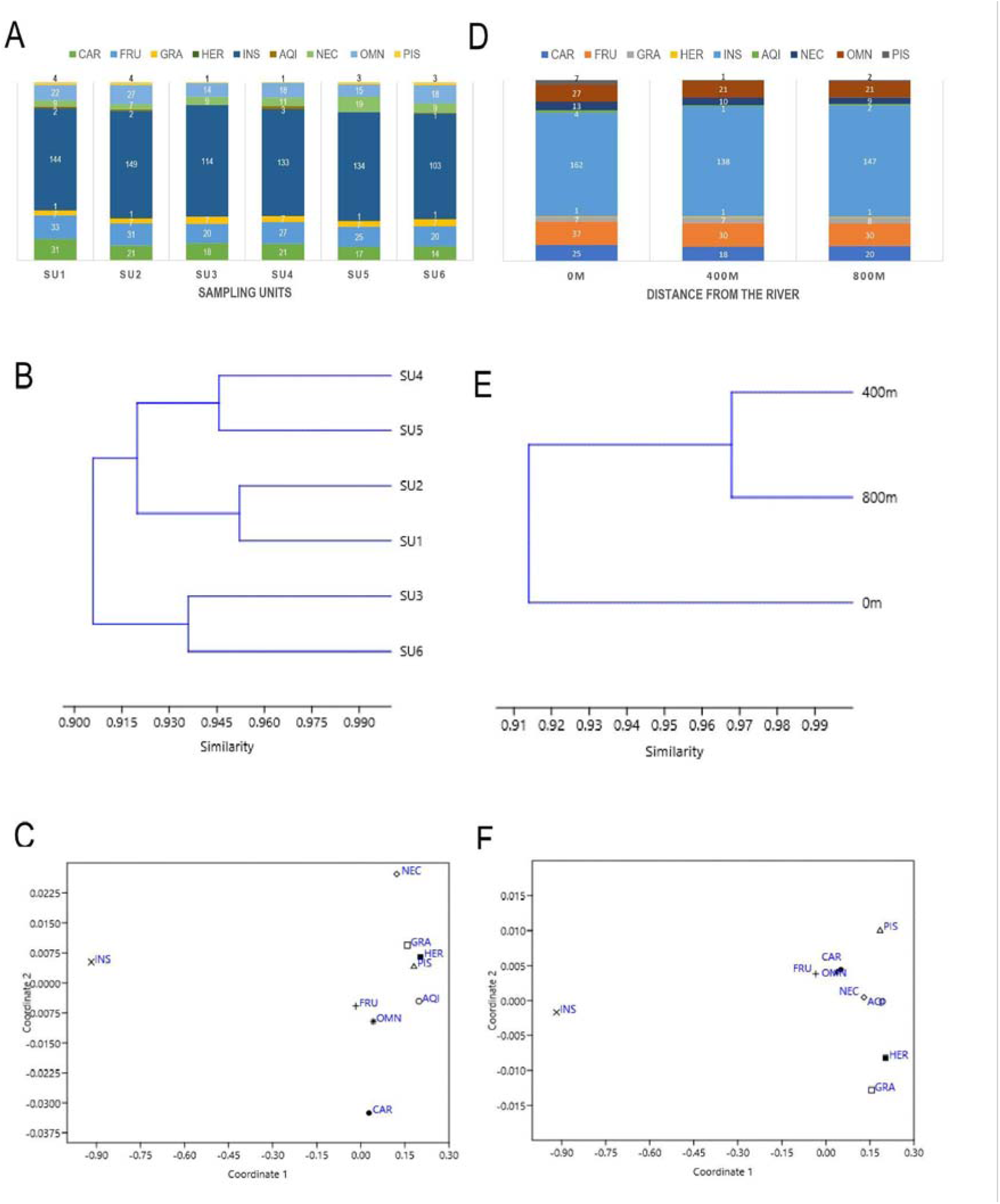
Distribution of bird food-web richness in the Kapawi River, Wayusentza community. (A) Distribution of the number of species by trophic groups in each sampling unit, (B) Cluster analysis based on the Bray-Curtis algorithm for the number of species per group in six sampling units, (C) MDS analysis for the data on the number of species per group in six sampling units; (D) Distribution of the number of species by trophic groups along the distance gradient, (E) Cluster analysis based on the Bray-Curtis algorithm for the number of species per group across three distance gradients, (F) MDS analysis of the data on the number of species per group in six sampling units across three distance gradients.

Across the sampling units, insectivores constituted the dominant functional group at all stations (x = 129 spp./order), with the highest species richness recorded in SU2 (*n* = 149 spp.) and SU1 (*n* = 144 spp.), coinciding with the sites of highest overall taxonomic richness. Frugivorous and insectivorous species exhibited a concerted numerical decline in SU3 and SU6, falling to their minimum values (Fig. 3A).

Key functional groups closely linked to the productivity of the water-land interface, such as specialists in aquatic invertebrates (Inv-Acua) and piscivores (Psc), showed a modest but persistent representation (x = 3 spp./ group), with peaks in species richness located in Sampling Units 1, 2, 4, and 5, and decreasing to the total absence of groups such as herbivores (Her) and aquatic invertebrates in SU3 (FIG. 3A). In contrast, granivorous species (Gra) demonstrated absolute and unchanging stability at exactly *n*= 7 spp./group in each and every one of the six units evaluated.

### Distribution of functional traits along the riparian distance gradient

When evaluating the vertical dimension of the riparian gradient (horizontal distance from the riverbank), it was found that the riverbank (0 m) exhibits the greatest species richness for nearly all of the ecological groups analyzed (Fig. 3B). Insectivores (*n* = 162 spp./group), frugivores (*n* = 37 spp./group), carnivores (*n* = 25 spp./group), and omnivores (*n* = 27 spp./group) reached their absolute maximums directly at the riparian interface. Groups with strict or facultative dependence on water resources, such as piscivores and aquatic invertebrate feeders, doubled and tripled their species richness at the water’s edge (*n* = 7 spp. and *n* = 4 spp., respectively) compared to the forest interior (400 m and 800 m). As the gradient moves inland (400 m), there is a general decline in the richness of nearly all ecological traits, followed by a slight selective recovery at 800 m, a pattern driven primarily by the return of insectivorous (*n* = 147 spp./group) and carnivorous (*n* = 20 spp./group) species. Herbivores remained perfectly stable (*n* = 1 spp./order) throughout the transect, while nectarivorous species (Nec) showed a marked affinity for the riparian zone (13 species at 0 m), stabilizing in the deep interior at a constant rate of nine species (Fig. 3B).

### Similarity Analysis and Functional Clustering

Similarity analysis based on the Bray-Curtis algorithm determined that the functional structure of the trophic groups maintains a high baseline homogeneity, with similarity exceeding 90% throughout the study (Fig 3B). The clustering dendrogram differentiated the network of stations into two main blocks: (a) a major node subdivided into two highly similar sub-clusters: the first comprising SU4 and SU5 (94.5% similarity), and the second closely associating SU2 and SU1 with a similarity level exceeding 95%; and (b) a clearly segregated secondary group linking SU3 and SU6 (93.5% similarity), which stand apart from the rest of the system below the 86% similarity threshold due to their lower levels of accumulated richness in the INS and FRU guilds.

Distance-based cluster analysis showed that the interior communities of the hardwood forest (400 m and 800 m) possess a highly coincident guild structure, sharing a similarity of nearly 97%. Meanwhile, the riparian community (0 m) is basally separated from the interior block, remaining isolated at an affinity scale of approximately 91.5% due to its high concentration of unique species and riparian specialists (Fig.3E).

The spatial ordering patterns obtained via MDS, both for the sampling units and along the distance gradient (Fig. 3C, F), confirmed the decisive influence of certain trophic traits on the structure of the avian community of the Kapawi River. In both two-dimensional projections (units and distances, Fig. 3C, F), Axis 1 exerts the primary control over biological variance, driven by a gradient of dietary specialization; thus, the following groupings are identified:

#### The Insectivore (INS) vector

In both analyses, the INS group is projected in a completely polarized and isolated manner at the negative left end of Axis 1. This demonstrates that the variation or loss of insectivorous species represents the dominant ecological force that alters the composition of the soundscape and destabilizes the homogeneity among the study sites.

#### The Frugivore (FRU) vector

It was positioned modestly but consistently toward the central negative region of Axis 1, acting as the second most influential group in the community structure.

#### The block of minority groups

The remaining groups (CAR, GRA, HER, AQI, NEC, OMN, and PIS) clustered very tightly at the far right (positive) end of Axis 1. This high graphical concentration indicates that, despite local variations in abundance, these trophic niches vary in a coordinated manner and maintain a proportionally stable abundance ratio throughout the entire monitored riparian corridor.

### Soundscape Analysis (Acoustic Indices)

Acoustic indices varied among recording stations and between species richness categories (Fig. 4). Overall, stations with higher species richness (≥250 spp) tended to exhibit slightly elevated values of ADI and AEI, suggesting greater acoustic diversity and a more even distribution of acoustic energy across frequency bands. In contrast, ACI showed relatively modest variation across stations, with no consistent separation between richness categories, indicating similar levels of temporal acoustic complexity (Fig. 5). The Bioacoustic Index (BI) displayed pronounced variability among stations, with higher values generally associated with high-richness sites, although with considerable overlap. NDSI values remained consistently high across all stations, reflecting a dominant biophonic component throughout the study area, with limited differentiation by richness category. Similarly, the Shannon index (SH) showed relatively stable distributions across stations, with only subtle increases in high-richness sites. Despite overlapping distributions, these patterns suggest that acoustic diversity (ADI) and evenness (AEI) are more sensitive to differences in species richness, whereas indices such as ACI and NDSI capture broader soundscape characteristics that are less strongly structured by local richness gradients.

**Figure 4.**
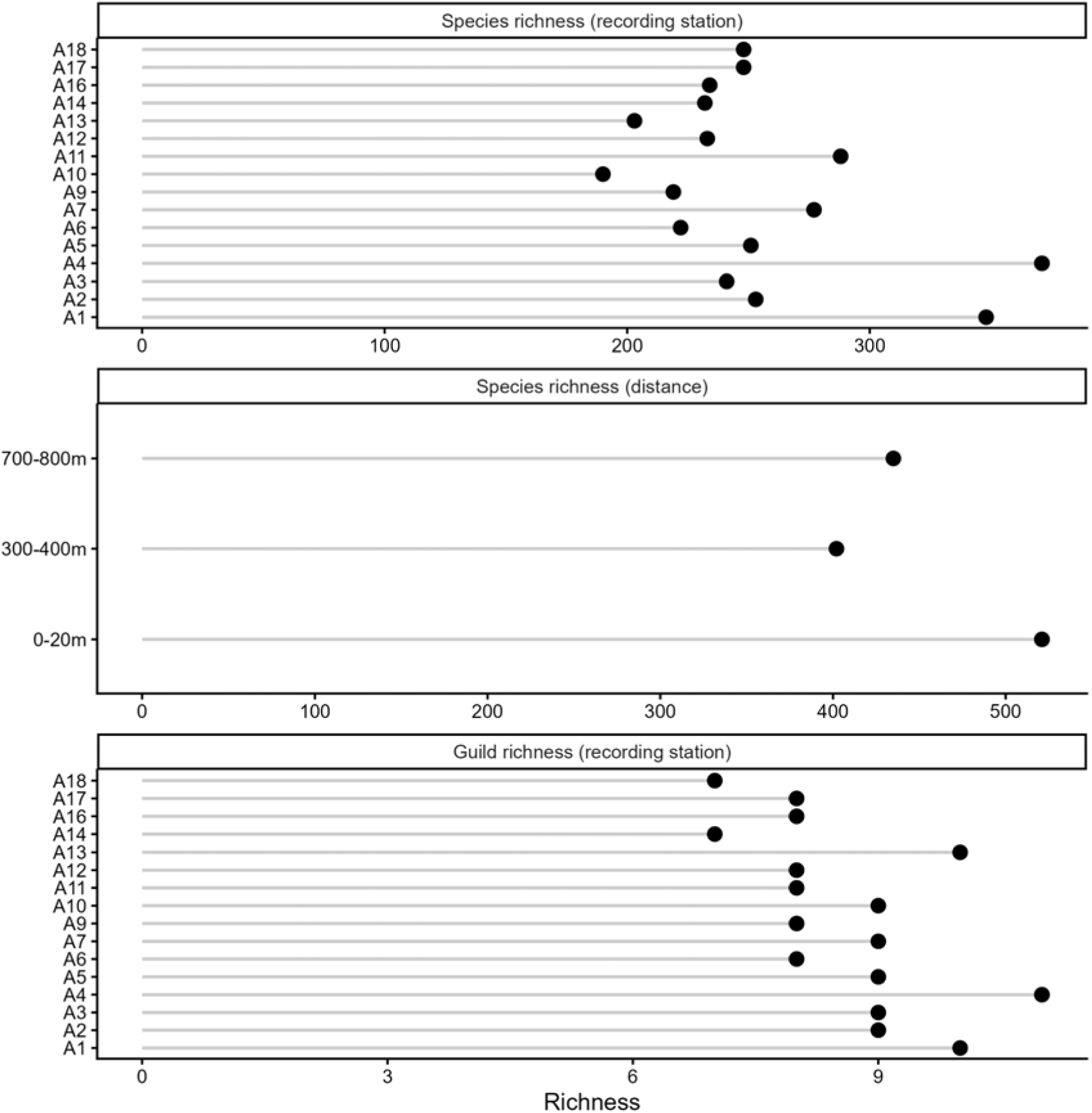
Acoustic-derived species and guild richness across recording stations and spatial locations.

**Figure 5.**
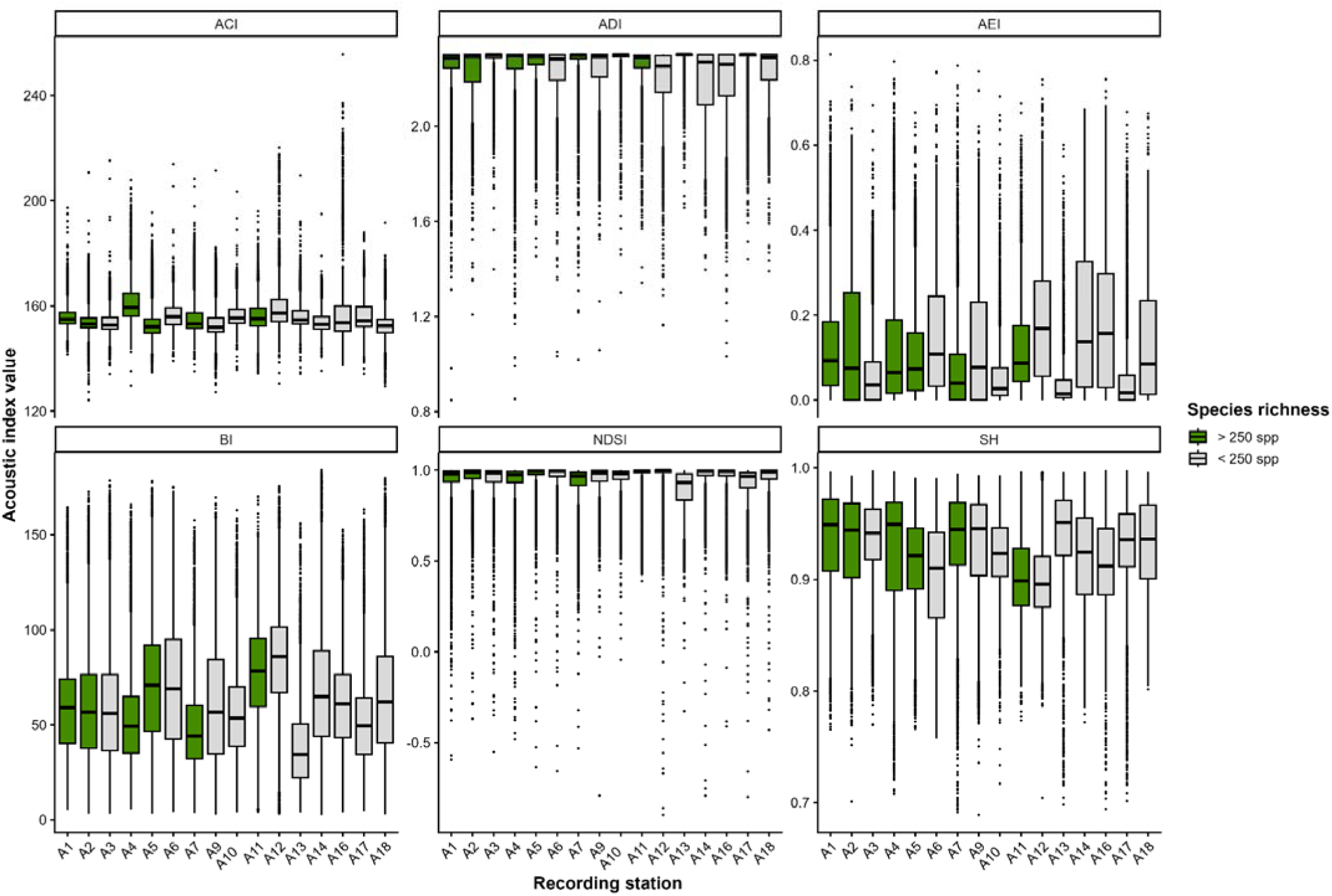
Comparison of acoustic indices among recording stations in the study area. SH = Spectral Entropy, NDSI = Normalized Difference Soundscape Index, ACI = Acoustic Complexity Index, ADI = Acoustic Diversity Index, AEI = Acoustic Evenness Index, BI = Bioacoustic Index.

Species richness showed distinct associations with the evaluated acoustic indices in a negative binomial GLM with a log-link function, indicating a structured response of the biological community to soundscape composition. Significant positive effects were observed for ACI (β = 0.0848, SE = 0.030, z = 2.79, p = 0.0053), ADI (β = 0.116, SE = 0.027, z = 4.29, p = 1.77 × 10), BI (β = 0.208, SE = 0.092, z = 2.27, p = 0.023), and SH (β = 0.115, SE = 0.056, z = 2.07, p = 0.039), suggesting that increases in acoustic heterogeneity, complexity, and structural diversity are associated with higher species richness (Fig. 6A, 6B). Among these predictors, BI showed the largest estimated coefficient on the model scale, followed by ADI, indicating comparatively stronger associations with species richness within the fitted model. ACI and SH, although of smaller magnitude, also exhibited significant positive effects, reinforcing the role of soundscape structure as an ecological predictor of biodiversity patterns. In contrast, AEI was not significantly associated with species richness (β = 0.0138, SE = 0.034, z = 0.402, p = 0.689), with estimates close to zero, suggesting a negligible contribution within the model. NDSI showed a negative but non-significant relationship (β = −0.0956, SE = 0.080, z = −1.19, p = 0.234), indicating a potential tendency for lower species richness in environments with higher relative contributions of anthropogenic sound; however, this pattern was not statistically supported. Model performance improved substantially relative to the null model, as indicated by a reduction in deviance from 60.19 to 15.74, reflecting a better fit of the model including acoustic predictors. The negative binomial formulation accounted for overdispersion in the response variable, providing an appropriate framework for modelling species richness. The model achieved an AIC of 161.17, supporting its adequacy for describing the relationship between bird species richness and soundscape structure.

**Figure 6.**
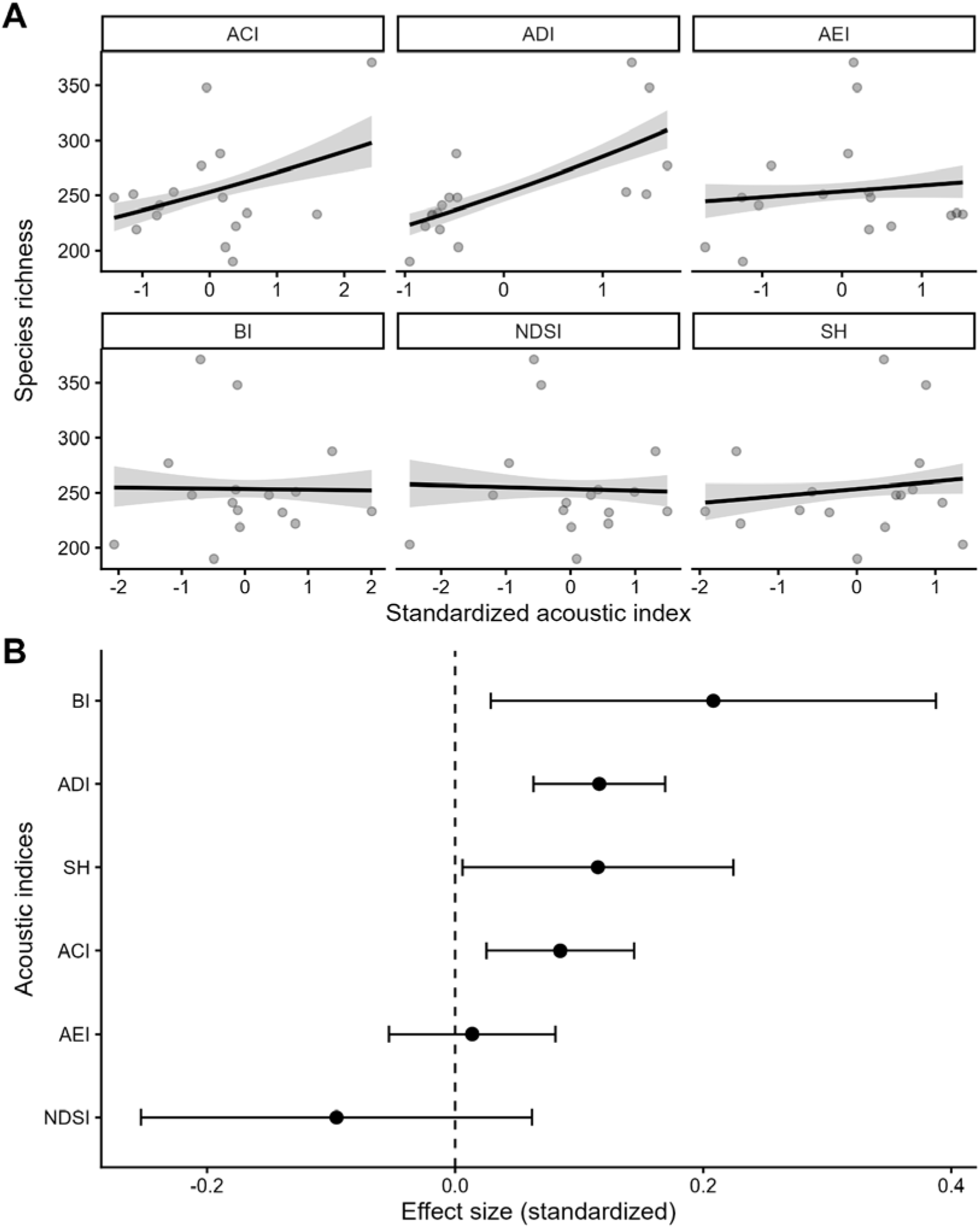
Acoustic indices predict species richness. **A.** GLM-fitted relationships between individual acoustic indices and species richness. Lines show model predictions. **B.** Standardized effect sizes (± s.e.) of acoustic indices derived from the GLM.

Acoustic indices exhibited distinct responses along the distance gradient from the river (Fig. 7). ACI showed relatively higher values at sites closest to the river (0–20 m), with a slight decline at intermediate and farthest distances, suggesting reduced temporal acoustic complexity inland. In contrast, ADI peaked at intermediate distances (300–400 m) and decreased toward the farthest sites, indicating that acoustic diversity is maximized away from the immediate river edge but not at the most distant locations.AEI displayed an increasing trend with distance, with the highest values observed at 700–800 m, reflecting a more even distribution of acoustic energy inland. Similarly, BI increased from near-river to intermediate distances, before slightly declining at the farthest sites, suggesting that overall biophonic activity is strongest away from the river margin. NDSI values were lowest near the river and highest at intermediate distances, indicating a relative reduction of biophonic dominance close to the river, likely due to increased geophonic noise (e.g., water flow). The Shannon index (SH) remained relatively stable across the gradient, with only minor variation among distances.

**Figure 7.**
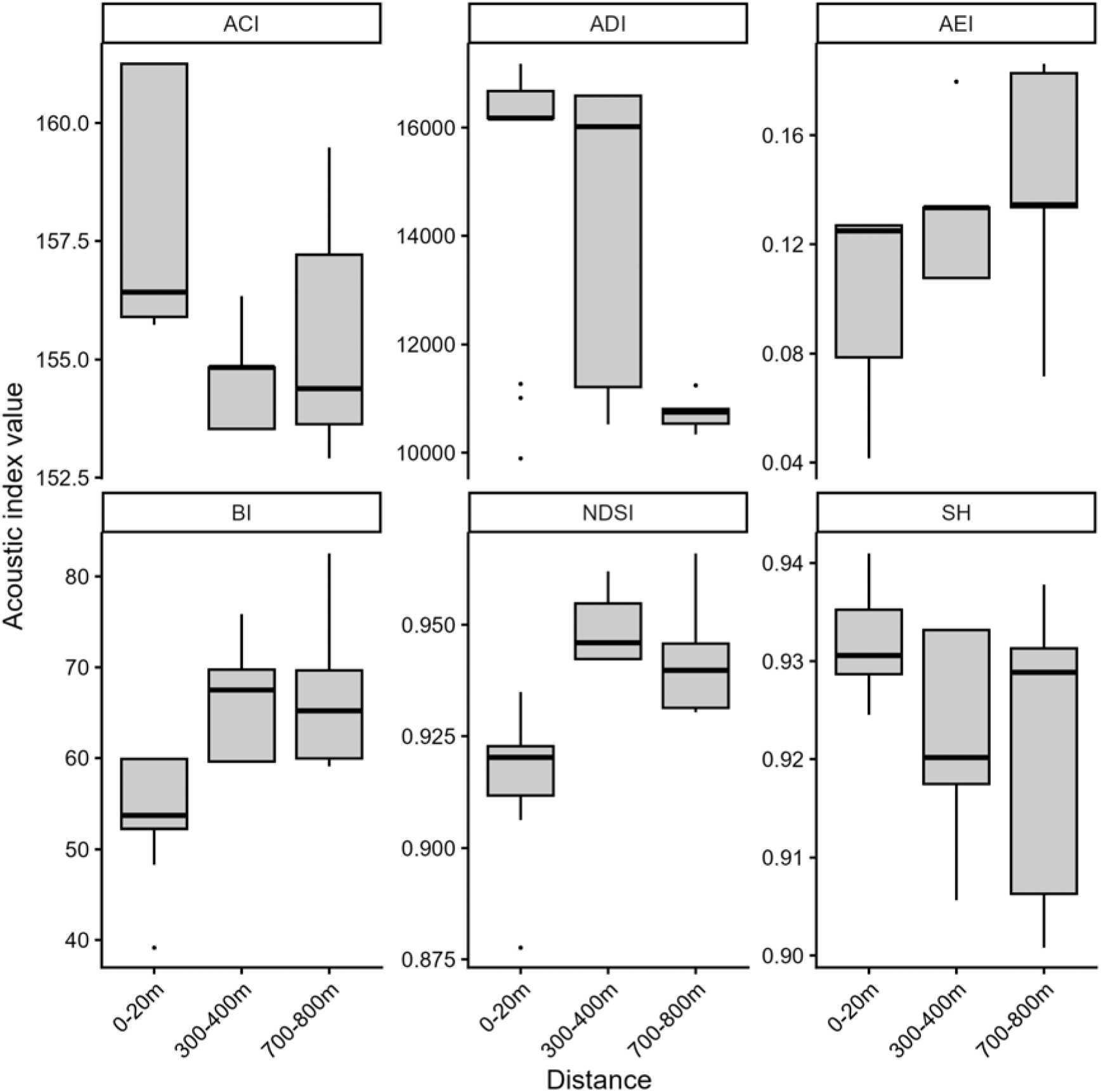
Variation in acoustic indices along the distance gradient.

The spatial distribution of acoustic indices along the distance gradient showed different patterns among metrics. NDSI values were relatively higher near intermediate distances, suggesting a balanced contribution of biophonic and geophonic signals, while lower values were observed toward the edges of the gradient (Fig. 8). In contrast, ACI and BI showed localized peaks, particularly in areas closer to the river and at distal inland points, indicating spatial heterogeneity in acoustic activity and sound complexity. ADI displayed a marked increase toward the farthest inland sites, suggesting higher acoustic diversity away from the immediate river influence. Similarly, AEI values were elevated in intermediate zones, reflecting a more even distribution of acoustic energy across frequency bands. The Shannon index (SH) remained relatively homogeneous across the landscape, with slight increases in mid-gradient areas, indicating moderate variability in acoustic diversity. Finally, these patterns suggest that proximity to the river influences the structure and distribution of acoustic communities, with distinct indices capturing different aspects of this spatial gradient.

**Figure 8.**
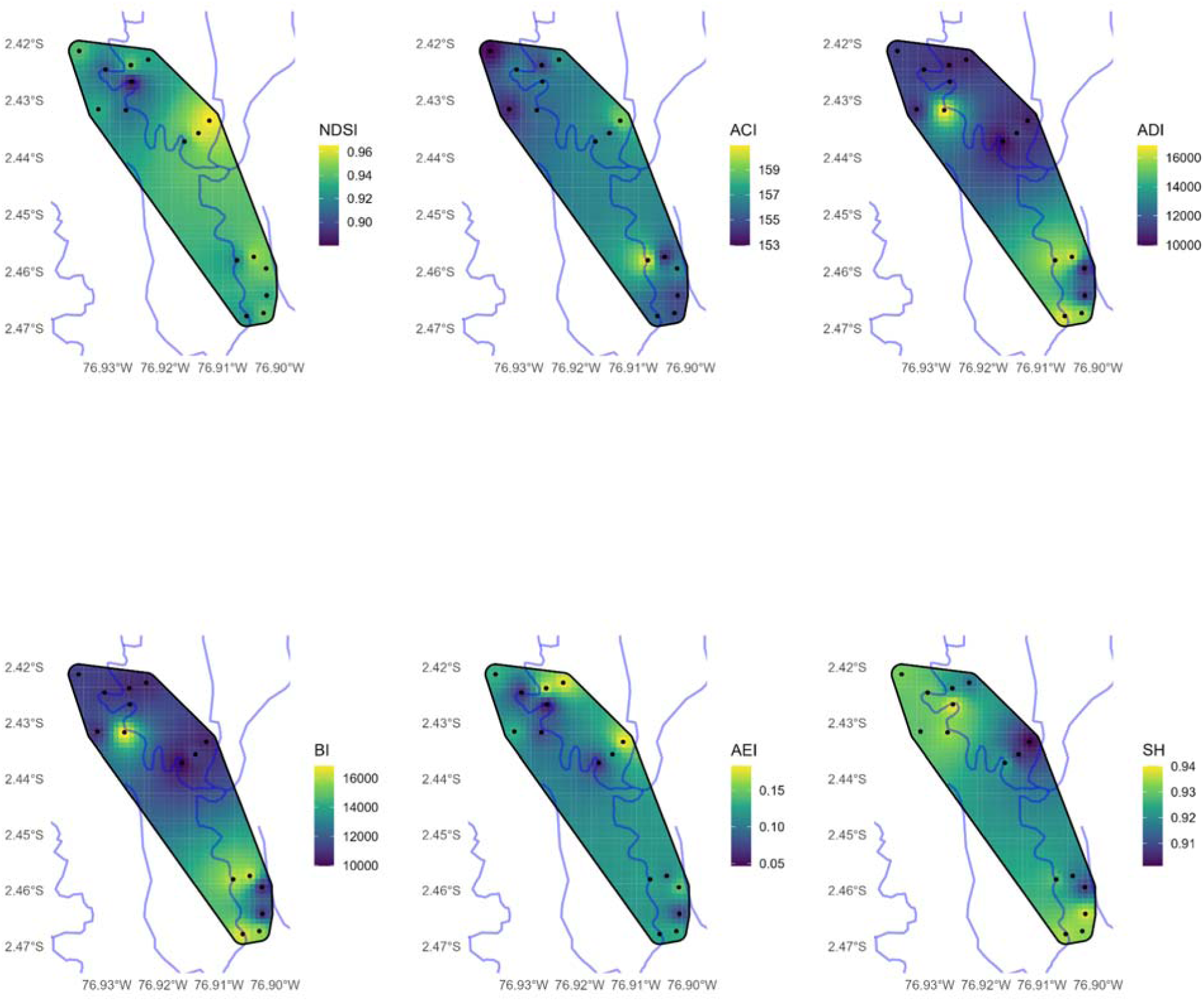
Variation in acoustic indices along the distance gradient, highlighting the main rivers within the study area. The Capahuari River is the closest hydrological feature to the recording stations.

## Discussion

Our data show that the Kapawi River corridor functions as a highly dynamic ecological system, where distance from the watercourse differentially modulates the taxonomic diversity and functional redundancy of bird communities. The findings confirm our hypothesis that the riverbank (0 m) acts as an ecotone of high productivity and biological uniqueness, recording the highest cumulative species richness (*n* = 325 spp.) and the highest number of endemic species (*n* = 71 spp.). This pattern of maximum biodiversity at the water-land interface is consistent with the findings of Naiman et al. (2005) and Remsen et al. (2020), who attribute the high turnover rate in Neotropical riparian zones to resource pulses and microclimatic gradients that generate specialized microhabitats.

Despite the pronounced differences in total species richness across the distance gradient, the analysis of variance (ANOVA) indicated that mean species richness per sampling unit did not differ significantly among distances. This phenomenon is explained by the high internal heterogeneity and local asymmetries along the riverbanks (ranging from *n* = 100 spp. to *n* = 209 spp.). In contrast, the forest interior (especially at 800 m) exhibited the greatest consistency in local species richness ( x = 151.0 ± 41.5 spp.). Non-metric multidimensional scaling (MDS) analyses for both species composition and functional traits (Fig. 2F, 3F) showed a polarized segregation of the riparian zone from the interior forest zones (400 m and 800 m). This marked hydrodynamic effect is consistent with the classic observations of Terborgh et al. (1990) in the western Amazon, where terra firme understory communities maintain a segregated structure isolated from riverine edge dynamics.

From a functional perspective, the structure of the dietary communities showed high baseline homogeneity, exceeding 90% similarity throughout the study, with an extreme affinity of 97% between the two inland sites (400 m and 800 m). The absolute dominance of insectivorous species (Ins), followed by frugivorous species (Fru), reflects the typical organization of mature lowland rainforests in the Neotropics (Bregman et al., 2016). MDS vector analysis determined that the loss or variation of insectivorous species is the dominant ecological force altering the composition of the soundscape. This is particularly relevant in the Amazon, where understory insectivores are classified as a group highly sensitive to subtle habitat modifications and acoustic stress (Sekercioglu et al., 2004). On the other hand, the restriction of dependent groups such as piscivores (Psc) and aquatic invertebrates (Inv-Acua) to the riverbank, along with the duplication of nectarivores (Nec), validates the existence of a strong niche partition driven by the structure of riparian vegetation.

The integration of automated bioacoustic tools using the GLM model demonstrated that the spatial structure of the soundscape accurately reflects these biological gradients. The significant positive effects of the ACI, ADI, BI, and SH indices indicate that sound complexity, heterogeneity, and diversity increase in tandem with bird richness. The Bioacoustic Index (BI) and the Acoustic Diversity Index (ADI) presented the most robust predictive coefficients, establishing themselves as excellent proxies for rapid biodiversity monitoring in Amazonian landscapes under indigenous management, in line with the suggestions of Ross et al. (2021) and Toledo et al. (2023).

Finally, the negative but non-significant relationship of the NDSI (β = **-**0.0956), combined with the lower biological richness detected in the units north of the community settlement (UM3 and UM6, associated with taxonomic contraction), suggests an emerging trend of riverine degradation or local disturbance in specific sections of the river. The frequency band analyzed for mechanical engines (1-2 kHz) is directly reflected in the IDW interpolation maps (Fig. 8), noting that, while the Kapawi corridor maintains outstanding ecological integrity, future increases in fossil-fuel-powered river transport could fragment the natural acoustic continuum, affecting the biological choruses essential for the communication and survival of Amazonian avifauna.

## Acknowledgments

To the Wayusentza community for their assistance in conducting the study.

## Notes

https://doi.org/10.6084/m9.figshare.32493456

https://doi.org/10.6084/m9.figshare.32494023

### Competing Interest Statement

The authors have declared no competing interest.

## References

Alcocer, I., Lima, H., Sugai, L. S. M., & Llusia, D. (2022). “Acoustic Indices as Proxies for Biodiversity: A Meta-Analysis.” Biological Reviews of the Cambridge Philosophical Society 97: 2209–2236.

Agisoft LLC. (2019). Agisoft Metashape Professional (Version 1.5.2) [Computer software]. Agisoft LLC.

Bregman, T. P., Lees, A. C., MacGregor, H. E., Darski, B., de Moura, N. G., Aleixo, A., Barlow, J., & Tobias, J. A. (2016). Using functional traits to assess the resilience of rainforest bird communities to tropical deforestation. Biological Conservation, 195, 258–267. 10.1016/j.biocon.2015.12.012

Boelman, N. T., Asner, G. P., Hart, P. J., & Martin, R. E. (2007). Multi trophic invasion resistance in Hawaii: bioacoustics, field surveys, and airborne remote sensing. Ecological Applications, 17(8), 2137–2144.

Depraetere, M., Pavoine, S., Jiguet, F., Gasc, A., Duvail, S., & Sueur, J. (2012). Monitoring animal diversity using acoustic indices: Implementation in a temperate woodland. Ecological Indicators, 13(1), 46–54.

Deichmann, J. L., Acevedo-Charry, O., Dey, L., Campos-Cerqueira, M., & Mitchell, A. (2017). It’s time to listen: Passive acoustic monitoring improves efficiency of tropical biodiversity surveys. Biotropica, 49(4), 482–489. 10.1111/btp.12434

Gibb, R., Browning, E., Glover-Kapfer, P., & Jones, K. E. (2019). “Emerging Opportunities and Challenges for Passive Acoustics in Ecological Assessment and Monitoring.” Methods in Ecology and Evolution 10:169–185.

Hammer Ø, Harper D, Ryan PD. 2001. Past: paleontological statistics software package for education and data analysis. Paleontologia Electron 4(1):9

Hill, A. P., Prince, P., Piña Covarrubias, E., Doncaster, C. P., Snaddon, J. L., & Rogers, A. (2018). AudioMoth: Evaluation of a smart open acoustic device for monitoring biodiversity and the environment. Methods in Ecology and Evolution, 9(5), 1199–1211.

Hilbe, J. M. (2011). Negative binomial regression. Cambridge University Press.

Isaaks, E. H., & Srivastava, R. M. (1989). Applied geostatistics. Oxford University Press.

Ismail, N., & Jemain, A. A. (2007). Handling overdispersion with negative binomial and generalized Poisson regression models. In Casualty Actuarial Society Forum (Vol. 2007, pp. 103–158). Citeseer.

Kahl, S., Wood, C. M., Eibl, M., & Klinck, H. (2021). BirdNET: A deep learning solution for avian diversity monitoring. Ecological Informatics, 61, 101236. 10.1016/j.ecoinf.2021.101236

Kasten, E. P., Gage, S. H., Fox, J., & Joo, W. (2012). The remote environmental assessment laboratory’s acoustic library: An archive for studying soundscape ecology. Ecological informatics, 12, 50–67.

Ministerio del Ambiente del Ecuador (MAE). 2013. Sistema de Clasificación de los Ecosistemas del Ecuador Continental. Subsecretaría de Patrimonio Natural, Quito, Ecuador

Naiman, R. J., Décamps, H., & McClain, M. E. (2005). Riparia: Ecology, conservation, and management of streamside communities. Elsevier Academic Press.

Pijanowski, B. C., Villanueva-Rivera, L. J., Dumyahn, S. L., Farina, A., Krause, B. L., Napoletano, B. M., … & Pieretti, N. (2011). Soundscape ecology: the science of sound in the landscape. BioScience, 61(3), 203–216.

Pieretti, N., Farina, A., & Morri, D. (2011). A new methodology to infer the singing activity of an avian community: The Acoustic Complexity Index (ACI). Ecological indicators, 11(3), 868–873.

Priyadarshani, N., Marsland, S., & Castro, I. (2018). Automated birdsong recognition in complex acoustic environments: A review. Journal of Avian Biology, 49(5), jav-01447. 10.1111/jav.01447

Remsen Jr, J. V., & Parker III, T. A. (1983). Contribution of river-created habitats to bird species richness in Amazonia. Biotropica, 223-231.

Remsen, J. V. (2020). The magnificent, enigmatic, and threatened avifauna of the Amazon basin. The Wilson Journal of Ornithology, 132(1), 1–12. 10.1676/1559-4491-132.1.1

Robinson, W. D., Hallman, T. A., & Betts, M. G. (2018). Can data-intensive bioacoustics provide accurate estimates of tropical bird abundance? Ecological Applications, 28(6), 1639–1648. 10.1002/eap.1757

R Core Team. (2025). R: A language and environment for statistical computing. R Foundation for Statistical Computing. https://www.R-project.org/

Ross, S. R. J., Friedman, N. R., Yoshimura, M., Yoshida, T., Donohue, I., & Economo, E. P. (2021). Utility of acoustic indices for ecological monitoring in complex sonic environments. Ecological Indicators, 121, 107114. 10.1016/j.ecolind.2020.107114

Sekercioğlu, Ç. H., Daily, G. C., & Ehrlich, P. R. (2004). Ecosystem consequences of bird declines. Proceedings of the National Academy of Sciences, 101(52), 18042–18047. 10.1073/pnas.0408049101

Sueur, J., Pavoine, S., Hamerlynck, O., & Duvail, S. (2008). Rapid acoustic survey for biodiversity appraisal. Biological Conservation.

Sugai, L. S. M., Silva, T. S. F., Ribeiro, J. W., & Llusia, D. (2019). Terrestrial passive acoustic monitoring: Review and perspectives. BioScience, 69(4), 273–286. 10.1093/biosci/biz011

Terborgh, J., Robinson, S. K., Parker, T. A., Munves, C. A., & Pierpont, N. (2021). Structure and organization of an Amazonian forest bird community. Ecological Monographs, 60(2), 213–238. 10.2307/1943045

Toledo, M. T., Campos-Cerqueira, M., & Aide, T. M. (2023). Moving beyond taxonomic lists: The role of passive acoustic monitoring in evaluating functional diversity in changing landscapes. Frontiers in Ecology and Evolution, 11, 1084321. 10.3389/fevo.2023.1084321

Ulloa, J. S., Aubin, T., Llusia, D., Bouveyron, C., & Sueur, J. (2024). scikit-maad: An open-source and modular Python toolbox for quantitative soundscape analysis. Methods in Ecology and Evolution, 15(2), 340–348. 10.1111/2041-210X.14201

van Merriënboer, B., Dumoulin, V., Hamer, J., Harrell, L., Burns, A., & Denton, T. (2025). Perch 2.0: The Bittern Lesson for Bioacoustics. arXiv. https://arxiv.org/abs/2508.04665

Wood, C. M., Kahl, S., Rahaman, A., & Zuckerberg, B. (2021). The machine learning revolution in bioacoustics: Optimizing convolutional neural networks for avian species identification. Global Ecology and Conservation, 31, e01875. 10.1016/j.gecco.2021.e01875

Wong, D. W. (2016). Interpolation: Inverse-distance weighting. International Encyclopedia of Geography: People, the Earth, Environment and Technology, 1–7.

